# Sustaining Seagrass Restoration: Long-Term Preservation of *Posidonia oceanica* Seeds and Seedlings

**DOI:** 10.1101/2025.03.18.643525

**Authors:** Alberto Sutera, Patrizia Spinelli, Davide Pacifico, Francesco Carimi, Roberto De Michele

**Affiliations:** Institute of Biosciences and Bioresources (IBBR), Italian National Research Council (CNR), Via Ugo la Malfa

## Abstract

Seagrass restoration efforts using *Posidonia oceanica* traditionally rely on lateral cuttings, a costly and labor-intensive method that damages parent meadows and limits genetic diversity. Seedling-based propagation presents a viable alternative which ensures genetic variability but is constrained by the unpredictability of seed stranding events and the short window for seed collection. To address this limitation, we tested methods for long-term seed and seedling storage, aiming to extend transplanting opportunities beyond natural germination cycles. We evaluated the effects of light or dark conditions, density, and seedling age on viability during storage at 4°C for six months. Our results demonstrate that light is essential for preserving viability, as all dark-stored samples died post-storage. Older seedlings (1-2 months old) exhibited higher survival rates (70-90%) compared to freshly collected seeds (40%). Seedling density did not significantly affect viability, simplifying large-scale storage. Attempts to induce dormancy with ABA or paclobutrazol were unsuccessful. This study provides the first successful protocol for seed and seedling storage, enabling year-round planning of seagrass restoration projects and enhancing their feasibility and efficiency.

## Introduction

*Posidonia oceanica* (L.) Delile is a species of seagrass endemic to the Mediterranean Sea, where it forms extensive meadows providing essential ecosystem services such as nursery grounds for fish and invertebrates, sediment stabilization, biodiversity enhancement, and carbon sequestration (Scanu et al., 2022). However, threats such as coastal development, pollution, and climate change have led to significant declines in its populations, necessitating effective conservation and restoration strategies (Telesca et al., 2015). Most restoration programs rely on clonal propagation by cuttings of lateral rhizomes from adult plants (Alagna et al., 2019). This approach further damages donor meadows, whose growth is very slow with a lateral expansion estimated at only 2 cm per year (Marbà & Duarte, 1998). Additionally, transplantation is laborious and costly since it requires specialized personnel for underwater operations. Finally, all transplanted cuttings are clones of the parent plants, which could affect the resilience of future meadows to environmental stressors and emerging diseases due to reduced genetic diversity (Sullivan et al., 2018). An alternative strategy relies on seedling transplantation, obtained by germinating seeds (Escandell-Westcott et al., 2023). *P. oceanica* reproduces sexually by releasing, in late spring, a large number of seeds, enclosed in a fleshy, buoyant fruit that allows dispersion along the prevailing currents. When currents and winds are unfavorable, a large proportion of fruits and seeds are washed ashore and can be conveniently collected by unskilled personnel while at the same time reducing the costs of collecting the propagation material. However, a propagation strategy based on seedlings is limited on the stochastic occurrence of stranding events, and on their short time frame, occurring on late spring (Belzunce et al., 2008; Sutera et al., 2024). Another major impediment is the lack of efficient protocols for the conservation of seedlings.

Expanding the germination period of *P. oceanica* seeds would offer several advantages to propagation efforts through its seedlings. First, it would allow to align the seedling developmental progress to the desires of the project, for example by reducing the number of seeds that are allowed to germinate at any given time, to optimize the management cost of dedicated facility and personnel. Second, it would present the opportunity to produce seedlings at the desired developmental stage at the optimal moment of transplantation, usually in spring and summer. Third, it would facilitate the transport of seeds and young seedlings between facilities for an extended period of time, due to the reduced dimension and fragility, compared to older seedlings and adult plants.

Many species of seagrass, such as *Zostera marina*, produce seeds that can remain dormant for extended periods, allowing them to survive in an unfavorable environmental condition. However, seeds of *P. oceanica* are not dormant, and germinate as soon as their fruits release them (Orth et al., 2000). The lack of natural dormancy makes the conservation of viable, ungerminated seeds a challenging task. Early attempts of storing *P. oceanica* seeds in the cold were unsuccessful, with their viability completely lost after just a few days of storage (Belzunce et al., 2008).

This article presents a comprehensive investigation into the factors affecting the viability and germination of *P. oceanica* seeds under controlled conditions, focusing on the influence of temperature, density, hormones, and light availability. Cold temperatures, while potentially detrimental to germination, can also extend the longevity of seeds in storage, providing a window of opportunity for future restoration efforts. By investigating various storage parameters and their effects on seed physiology, we seek to identify optimal conditions that preserve seed viability over extended periods. Our findings will contribute to the development of best practices for seed conservation and management, ultimately supporting the restoration of *P. oceanica* in degraded habitats.

## Materials and Methods

### Seedling Storage Experiment

The seeds of *P. oceanica* stranded onshore were collected on May 19, 2022 on the beach of Trabia (37.994 N, 13.666 E) and on May 31, 2024 on the beach of Mazara del Vallo (37.662 N, 12.542 E), both located in northwest Sicily, along the Tyrrhenian coast (Fig. S1 A-B). We only collected fresh seeds by scanning the shoreline during the incoming tide period. Dry seeds and fruits, and those found away from the crashing waves, were discarded. The seeds were washed with seawater to remove debris, brought to the laboratory and stored overnight at 10°C in a bucket with 15 L of artificial seawater (ASW, 3.8% salinity). For each collection event, seeds were then transferred to two aquaria in batches of 300 seeds, with 40 L of ASW, equipped with pump, filtration, growth light (300 lux with a photoperiod of 14h/10h light/dark) and the water temperature set at 18°C (Fig. S1 C-E). Seeds were inspected daily and those showing early symptoms of rot and/or mold were promptly removed. The water was changed weekly and the filters monthly. At the experimental time points (0, 4 weeks, 8 weeks), batches of 50 seeds/seedlings, randomly picked, were photographed, their growth measured (maximum leaf length) and transferred to a fridge at 4°C. The variables tested for storage were the following:

1) age of the seedlings: freshly collected seeds (T0) or seedlings that had already developed in the aquaria for one (T1) or two (T2) months.
2) light conditions: cold storage in either dark or under illumination by grow light (300 lux with a photoperiod of 8h/16h light/dark);
3) Density: mass storage of the whole batches in transparent plastic beakers with 2.5 L ASW, or storage of individual seedlings in glass culture tubes (20 cm tall, 2 cm wide) with metal lid, filled with 25 mL ASW, arranged in racks (Fig. S1 F-G).
4) Hormonal treatment: only for freshly collected seeds stored in the dark in individual tubes, we treated with the dormancy-inducing hormone abscisic acid (ABA) or paclobutrazol, an inhibitor of the germination-promoting hormone gibberellic acid. The treatments were carried out in batches of 10 seeds per treatment. Both ABA and paclobutrazol were prepared as 30 mM stock solution in ethanol, and provided at a final concentration of 10 µM, 100 µM (ABA), and 10 µM, 100 µM, 1 mM (paclobutrazol) directly in the storage ASW. Control seeds were treated with 833 µL ethanol, the solvent present in the highest dose of treatment. Treatments were freshly provided at each ASW change.

ASW was replaced every week for the first month, then monthly. At each water change, the seeds were inspected and those that appeared to rot were removed. Two batches of 50 seeds were left in two different aquaria as a growth and viability control. Each month, dead and rot seedlings were removed and the other seedlings photographed and their maximum leaf length was measured. Six months after seed collection, all samples were moved to the aquaria with the same initial conditions. After further two months in aquaria, we evaluated viability. Leaf growth was measured monthly up to three months after transfer to aquaria.

### Viability and growth measurements and statistical analyses

Seeds and seedlings appearing soft, breaking apart upon handling, and/or covered by mold were discarded and counted as rot. The remaining samples, during the cold storage period, were considered firm and potentially viable. Viability was eventually scored after two months of growth in aquaria. Seedlings that had not developed from seeds, that appeared rot, molded, brown or that had not grown at all, were considered dead.

Leaf growth was measured in photographs to minimize handling of seedlings and permanence in dry conditions. The maximum leaf length was measured by ImageJ processing, using a ruler as a scale.

Each experimental batch, as described above, was replicated in the two seed collection events. Data are presented as aggregated for leaf length and as mean and standard deviation for firmness and viability. The statistical significance of viability comparisons was tested by pairwise Chi-square tests of independence, since it is a binary category, with α = 0.05. Leaf lengths had unequal distribution sizes among treatments, due to the different numbers of surviving seedlings, and therefore were analyzed by Kruskal-Wallis test followed by pairwise Mann-Whitney tests, with α = 0.05.

## Results

Untreated, control seeds in aquaria developed into seedlings. Mortality increased sharply in the first six weeks, when approximately 23% of the seeds had covered in whitish mold and eventually rotted (Fig. 1A-E). Mortality increased again, at a less pronounced pace, after four months of growth in the aquaria, eventually reaching 50% after 9 months. The surviving seedlings grew at a constant rate, with leaves elongating up to 142 ± 41 mm after nine months in the aquaria (Fig. 1F-G).

**Fig. 1:**
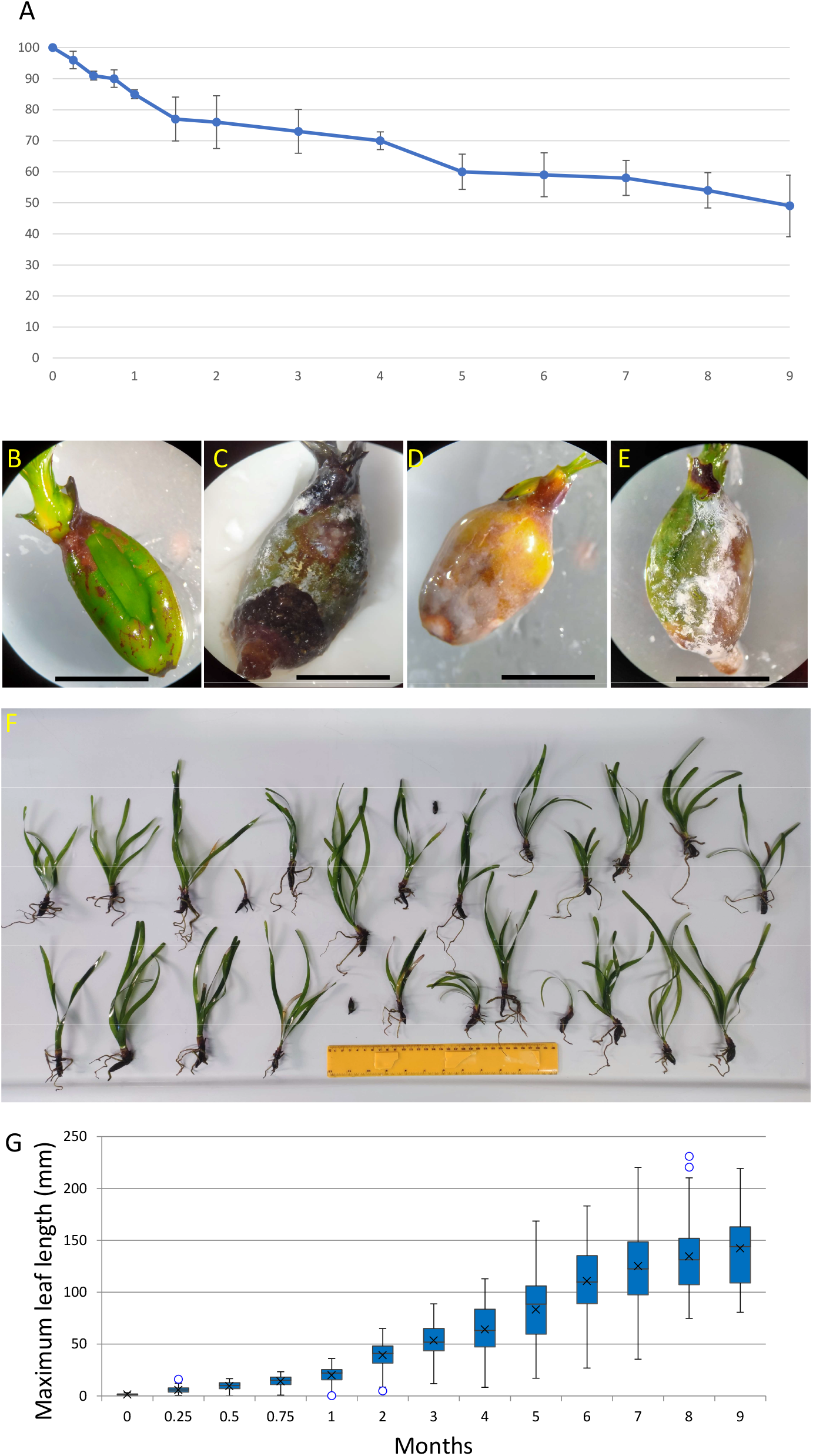
Development of *P. oceanica* seedlings in aquaria. Viability of seedlings over time (A). Representative images of a two-weeks old healthy seedling (B) and different examples of dead and rotting seedlings (C-E), scale = 1 cm. Appearance of seedlings after nine months of growth (F). Seedling growth over time, measured as maximum leaf length (G).

To develop protocols for long term conservation of *P. oceanica* seedlings, we measured firmness and growth up to six months after storage in the cold, either in the dark or under moderate light conditions, and with seedlings individually preserved in tubes or kept together in containers (Fig. S1 F-G). We also tested whether the seedling stage of development had an effect on preservation, by storing freshly collected seeds (T0), 1-month old seedlings (T1), and 2-months old seedlings (T2).

The effect of cold storage on rotting depended on the age of seeds/seedlings and on the light conditions. In the light, fresh seeds (T0) were most affected by prolonged cold storage (Fig. 2A). Over time, many of them progressively rotted and were discarded. After six months of storage, only 67% and 81% of the seeds stored in lit beakers and tubes, respectively, maintained firm and were transferred to the aquaria for testing development. Germinated seedlings were more resistant to storage conditions under light treatment, especially in beakers. At the end of the cold storage period all 1- and 2-months old seedlings (T1 and T2) kept in lit bakers were firm. For seedlings stored in individual tubes, 95% of 1-month old seedlings (T1) were firm at the end of the cold storage period. All older seedlings (T2) in tubes maintained firm. In the dark, all seeds and seedlings maintained firm up to six months, regardless of their age and container type. However, the seedlings stored in the dark presented darker leaves and often brown tips, while the leaves of seedlings stored in the light condition were homogeneously green (Fig. 2B).

**Fig. 2:**
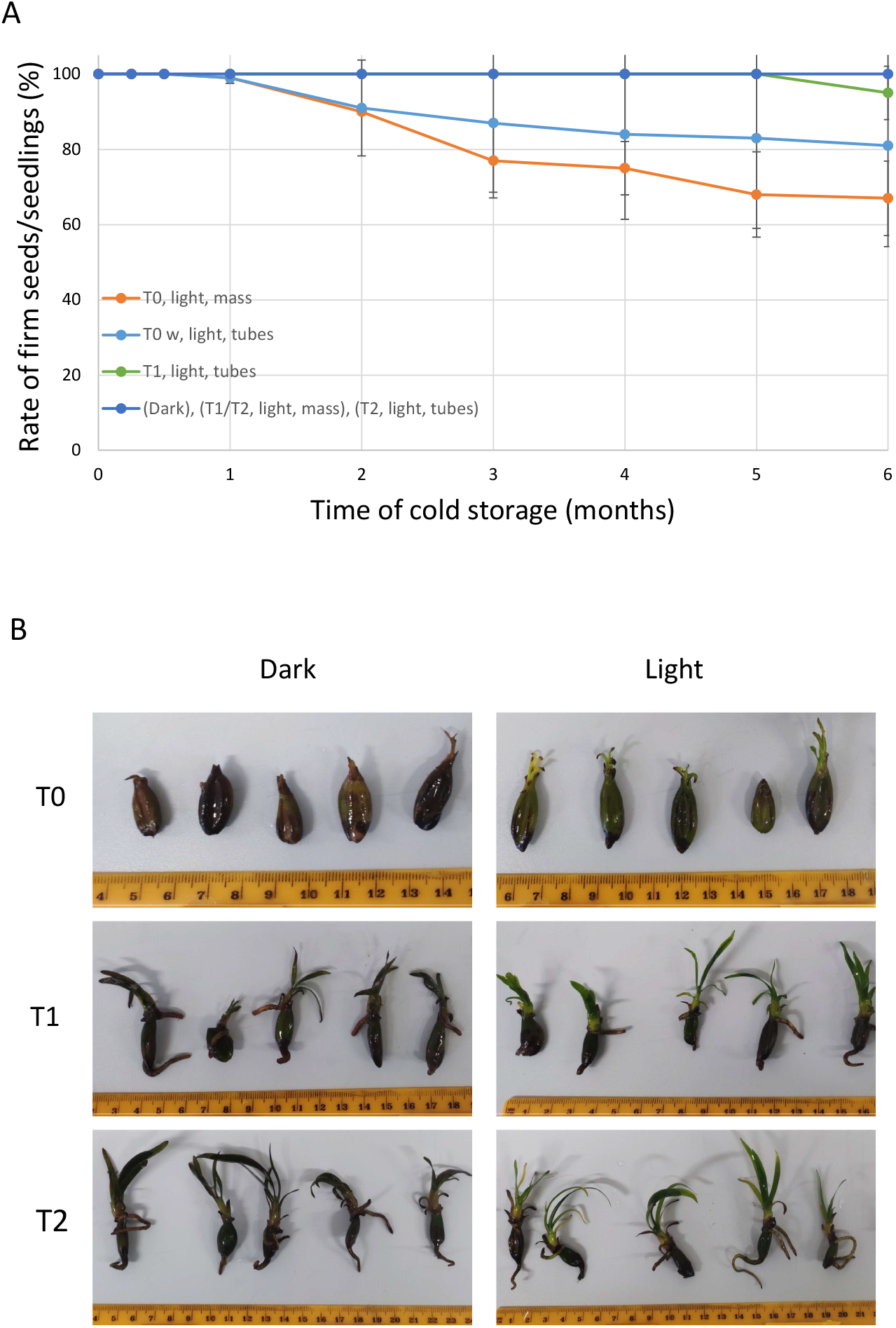
Proportion of firm seeds and seedlings stored at 4°C over time. Mean ± standard deviation (A). Representative images of the seedlings at the end of the mass storage period in light and dark (B). Note the different scales of the images.

During the storage period in the cold, growth of seedlings was stopped or severely delayed (Fig. 3A-D). No significant differences in leaf growth were observed between seedlings stored in the dark or light conditions, or between container types.

**Fig. 3:**
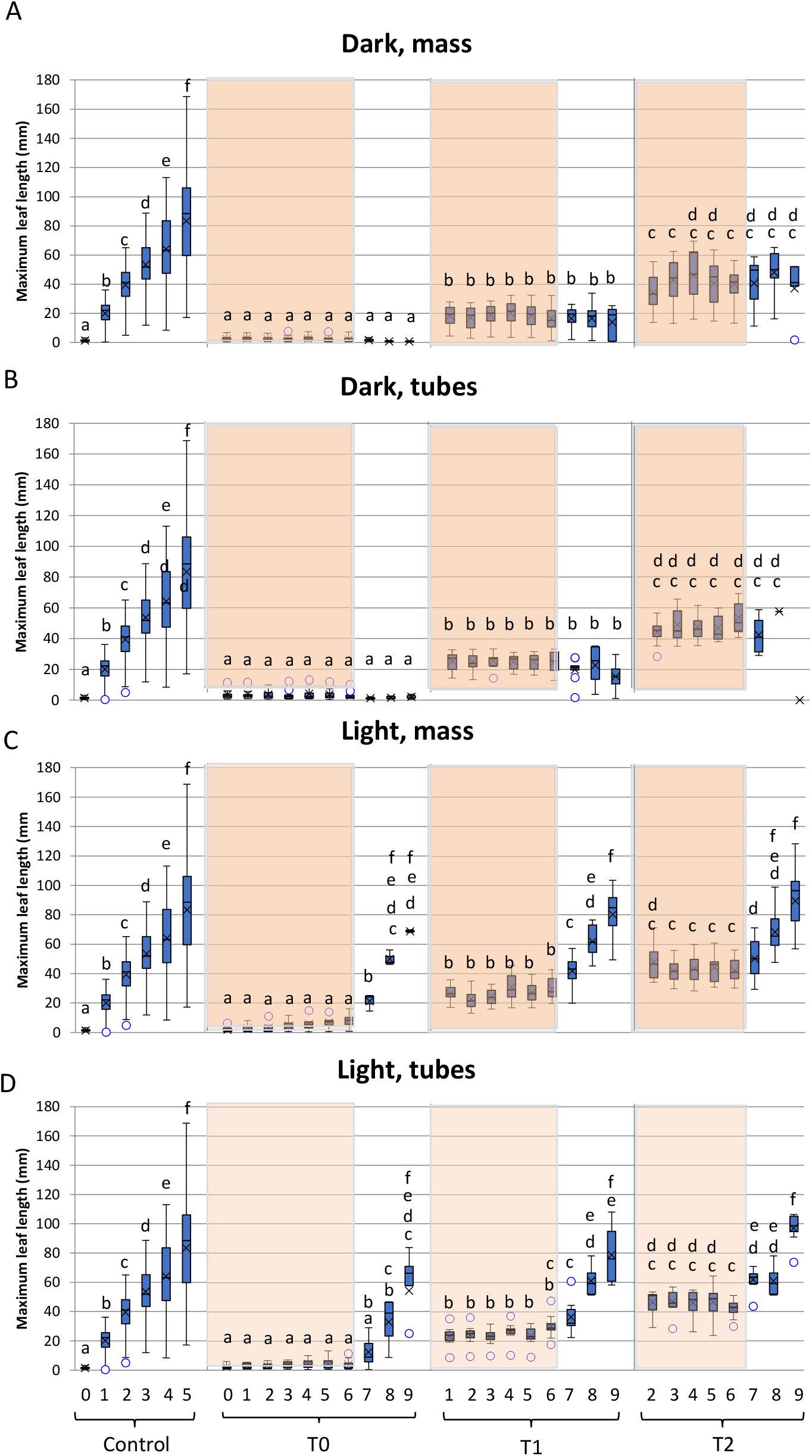
Seedling growth along time, measured as maximum leaf length, during storage and after transfer to aquaria. Seedlings stored in the dark in beakers (A); seedlings stored in the dark in tubes (B); seedlings stored in the light in beakers (C); seedlings stored in the light in tubes (D). Shaded areas indicate the time of storage in the cold; clear areas indicate time of active growth in the aquaria. Different letters indicate significant differences along time points among treatments.

After a storage time of six months at 4°C, we transferred the seedlings to aquaria with optimal growth conditions to monitor whether the storage treatment affected the potential to develop normally. After further two months in aquaria at 18°C, we counted the number of viable and dead seedlings for each treatment (Fig. 4A). Strikingly, all seeds and seedlings that had been stored in the dark died. Some of them just rotted, and the others, although firm, were dark brown and their leaves did not elongate (Fig. 3A-B). On the other hand, cold storage in the presence of moderate light conditions preserved the viability of most seedlings (Fig. 4A). Seeds that had been immediately stored after collection in beakers or individual tubes (T0) developed into seedlings at a final rate of 38% and 40%, respectively, including the fraction that had already rotten during the storage period. Higher survival rates (63-70%) were observed from seedlings that had been allowed to develop for one month before cold storage in beakers and tubes (T1), similar to control seedlings. Two-month old seedlings stored in the cold (T2) showed the highest survival rates, 89-90%. The differences in survival rates between the two types of containers were not significant for any class of seedlings.

**Fig. 4:**
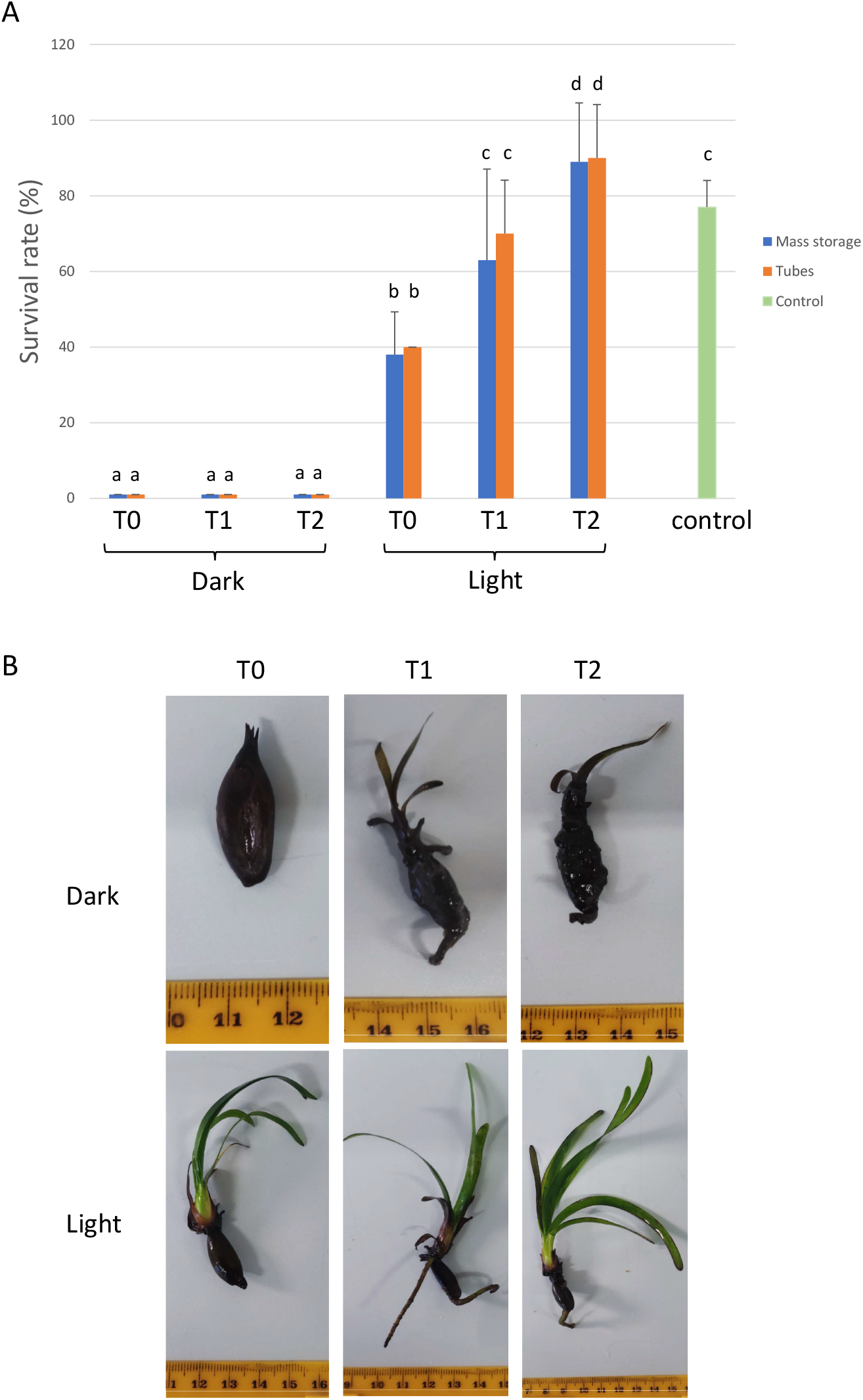
Viability of seedlings two months after transfer to aquaria. Survival rate (A). Value of control is relative to four months-old seedlings. Different letters indicate significant differences along time points among treatments. Representative seedlings at the end of the growth test (B). Note the different scales of the images.

Surviving seedlings in aquaria resumed growth in a fashion similar to control seedlings of the same age that had never been stored, as observed by the maximum leaf length (Fig. 3C-D, Fig. 4B). Seedlings appeared green and healthy with steady growth after transfer to aquaria up to three months, when we interrupted our measurements.

To test whether it was possible to induce dormancy in *P. oceanica* seeds and maintain their viability under cold storage in the dark, we treated freshly collected seeds with 10 µM,100 µM ABA or with 10 µM . 100 µM, 1 mM paclobutrazol, a GA inhibitor. After six months of cold storage at 4°C, the seeds were washed and transferred to aquaria at 18°C to test viability and growth. No seedlings developed from any seed, indicating that cold storage in the dark for six months was lethal even in the presence of dormancy-inducing hormonal treatments (data not shown).

## Discussion

The choice of the propagation material for seagrass restoration projects has practical, economic, and ecological consequences. For *P. oceanica*, restoration efforts traditionally employ lateral cuttings from parent meadows, assembled in an anchoring structure and fixed to the substrate. This strategy is cost- and labor-intensive, requiring specialized personnel operating underwater. Additionally, it damages the parent meadows and results in clonal replication, thus more susceptible to diseases and environmental variations. In contrast, a seedling-based propagation strategy presents several advantages. During stranding events, thousands of seeds are available for convenient collection by unskilled personnel, with no damage to existing meadows. Since seeds are the result of sexual reproduction, the genetic diversity of seedlings is preserved, potentially improving the resilience of transplanted populations. Despite the fact that no data are available on the viral load of *P. oceanica* seeds, in flowering plants seeds are also generally deprived from viruses present in mother tissues, further improving the fitness of seedlings compared to plants derived from cuttings (Bradamante et al., 2021). Developing seedlings produce adhesive roots, providing a strong attachment to the substrate, which can then be arranged on the seabed (Alagna et al., 2020).However, stranding is a punctual, unpredictable event, occurring only on a few days on late spring. Stranding is characterized by high spatial and temporal variability. Depending on daily winds and currents, stranding can occur on certain beaches and not others, despite the proximity of meadows. Predicting the days of maximal stranding is also challenging, since events can be scattered along several weeks. Furthermore, flowering and seed release do not occur every year, especially on the northern and colder Mediterranean shores, although events are becoming more and more frequent, probably due to climate change (Balestri et al., 2017; Diaz-Almela et al., 2007). In southern, warmer regions, such as the Sicilian coast, stranding events occur almost every year (Calvo et al., 2010).

Once stranded, seeds are exposed to dry sand, sunlight, and heat, seriously affecting their viability. Recent evidence suggests that the viability of stranded seeds is preserved for hours when seeds are enclosed in fruits or protected by direct sunlight. On the contrary, naked and sun-exposed seeds rapidly die (Sutera et al., 2024). The effective time for collection of hundreds of viable stranded seeds is therefore limited to a few hours after stranding, especially early in the morning or late in the afternoon. However, incoming tides usually retrieve part of stranded fruits and seeds, thus urging the collection process (Sutera et al., 2024).

In order to produce seedlings to be used as propagation material for restoration programs, collected seeds need to be rapidly placed in growing tanks, equipped with filtering systems and temperature control. In the first weeks of culture, typically 12-45% of seeds and young seedlings rot or die (Balestri et al., 1998; Belzunce et al., 2008; Buia & Mazzella, 1991; Sutera et al., 2024). In our experiments, we observed the same trend, with up to 23% of the seedlings rotting in the first six weeks. The causes of this massive mortality are unknown, but they may involve fungal and/or bacterial infections, as suggested by the presence of biofilms in the decaying material (Alagna et al., 2024). Removal of rotting seeds and replacement of water are essential in promoting seed viability in this stage of development. After the first weeks, the seedlings become more resistant and their mortality rate drops. Under our experimental conditions, seedling viability showed a second phase of modest decrease after four months, eventually reaching 50% after nine months. It is possible that at this stage the seedlings are already too large (mean leaf length about 15 cm) and too dense for our tanks. A second hypothesis for the decrease in viability is that, unlike the natural environment, seedlings in tanks were kept at constant photoperiod and temperature, eventually stressing the plants.

The handling of the large number of seedlings produced by stranding events poses management challenges due to the space and time constraints of the facility and the personnel involved in the restoration projects. Furthermore, the synchronized event of seed release sets a fixed calendar in seedling development, anchoring to the substrate, and transplant to the areas to be restored. Considering that the establishment of efficient root anchoring to the substrate requires at least five months (Alagna et al., 2015), and that prolonged storage in tanks extends the cost and effort of propagation project, due to density, increased size, and facility occupation and management, the optimal age for seedling transplant is about 6-7 months (Alagna et al., 2020; Zenone et al., 2025). However, considering that seed release usually occurs around mid-May, and that winter months can be particularly challenging for transplantation due to frequent storms, which affect both underwater operation and subsequent seedling colonization on unconsolidated substrate (Alagna et al., 2015; Infantes et al., n.d.; Zenone et al., 2022), there is a misalignment between optimal seedling age and the most convenient transplanting period. By setting a protocol for effective long-term conservation of *P. oceanica* seeds, our objective was to extend the period of seedling production and transplantation, uncoupling them from the natural calendar of seed germination. Since *P. oceanica seeds*, unlike other seagrasses, are not dormant, we tested temperature, density, light conditions, and hormonal control as factors affecting viability during long-term storage.

Our results show that light is the most important factor in preserving seed and seedling viability during storage in the cold. All samples stored in the dark, regardless of their age and density, died at the end of the storage period, as shown by their aspect and the inability to grow further once they were transferred to growth conditions. Our data agree with what observed in (Belzunce et al., 2008), where dark storage at 5°C for just 11-17 days was sufficient to suppress viability. Contrary to what was hypothesized in that paper, however, cold storage is not the factor that affects seed viability, since we observed that seeds stored in the the cold in the presence of light were viable up to six months. Germinated seedlings can survive up to three months in the shade, mobilizing carbohydrate reserves from seeds (Celdran, 2013). It remains to be tested whether cold storage periods in the dark shorter than six months can be tolerated by germinated seedlings. Interestingly, dark stored seeds and seedlings maintained firm throughout the storage months, more so than samples stored in the light. A possible hypothesis is that in the light viable seedlings exudate molecules capable of promoting the microbial community (bacteria and fungi) affecting weak seedlings, which eventually rot. In the dark, such metabolic activation would be prevented and rotting occurred only once dead seedlings were transferred to illuminated and warmer conditions. In agreement with this hypothesis, seeds stored in the dark do not mobilize their carbohydrate reservoirs (Celdran, 2013).

A large portion of light-stored seedlings maintained viable and resumed growth once transferred to aquaria, with rates similar to those of control plants. For long-term storage in the cold, therefore, light represents an essential condition to preserve viability. Light clearly promotes photosynthesis, which is apparently required for such long storage. Interestingly, it is known that seeds, the only organ present in T0 samples, are photosynthetically active in *P. oceanica*. Furthermore, even in three months-old seedlings, the rate of photosynthesis by seeds exceeds leaves photosynthesis by 4-5 times (Celdran, 2013). However, cold temperature prevents active growth, since we did not observe any leaf elongation in light-exposed stored seedlings.

In addition to light, the stage of seedling development was the factor that most influenced viability during storage. Freshly collected seeds were more affected by storage, and less than 40% of seeds eventually developed into healthy plants once growing conditions were resumed. Older seedlings were more resistant, with survival rates reaching 70% for 1-month old seedlings and even 90% for 2-months old seedlings. Interestingly, the viability of oldest stored seedlings exceeded that of the control plants of the same age (around 70%). However, this figure is not totally unexpected, since viability of control plants suffered from the sharp initial decrease we described earlier, probably due to damaged, weak, infected or otherwise ‘doomed’ seeds. The T1 and T2 seedlings, on the other hand, were selected from the pool among the surviving seedlings, so they already belonged to the resistant fraction of the plant population. As a recommendation, we endorse the storage of seedlings at least 1-month old, rather than freshly collected seeds. However, the extra growth time prior to storage imposes a layer of complexity in the protocol management, which might affect the seedling production pipeline at larger scales. Therefore, in each project, a trade-off between viability preservation and handling steps must be addressed. In some cases, the final 40% viability score could be sufficient to justify the immediate storage of fresh seeds, especially considering that about 30% of seeds would inevitably die even in control populations under ideal growing conditions.

The seedling density did not affect viability for any age group. Considering the additional work burden in maintaining seedlings in separate vials, especially during water changes, and the increased space requirements of tube racks compared to beakers, it is conveniently much easier to store seedlings together in batches in large enough containers. Next protocol optimization might focus on maximizing the seedling density by testing different water volumes, container types, and number of seedlings per batch.

Finally, we were unable to artificially induce dormancy in *P. oceanica* seeds, by providing the dormancy-inducing hormone ABA or paclobutrazol, an inhibitor of the germination-inducing hormone GA, at concentrations that proved useful in other studies (Goggin et al., 2009; Khan, 1994; Nakabayashi et al., 2012; Steinbach et al., 1997). It is possible that the use of these modulators at shorter storage times in the dark or in the copresence of light might enhance the protocol of seed conservation. However, considering the cost of these compounds and the additional handling operation required, it is probably not worth investing in this strategy for large-scale seedling production projects.

We limited our storage test to six months after seed collection, in order to cover a full year of seedling availability until next stranding events, if we consider that usually transplanted seedlings are at least six months old. Most likely, shorter storage times result in increased viability, possibly even for samples stored in the dark. Future tests might assess these timings.

In summary, this is the first study to report successful protocols for the storage of *P. oceanica* seeds and seedlings, preserving their viability and ability to develop normally into healthy plantlets, de facto uncoupling seed stranding events from seedling development. Reforestation projects now have the possibility of planning plant production and transplants year round, with evident economic and practical advantages for their implementation.

## Figure legends

**Fig. S1:**
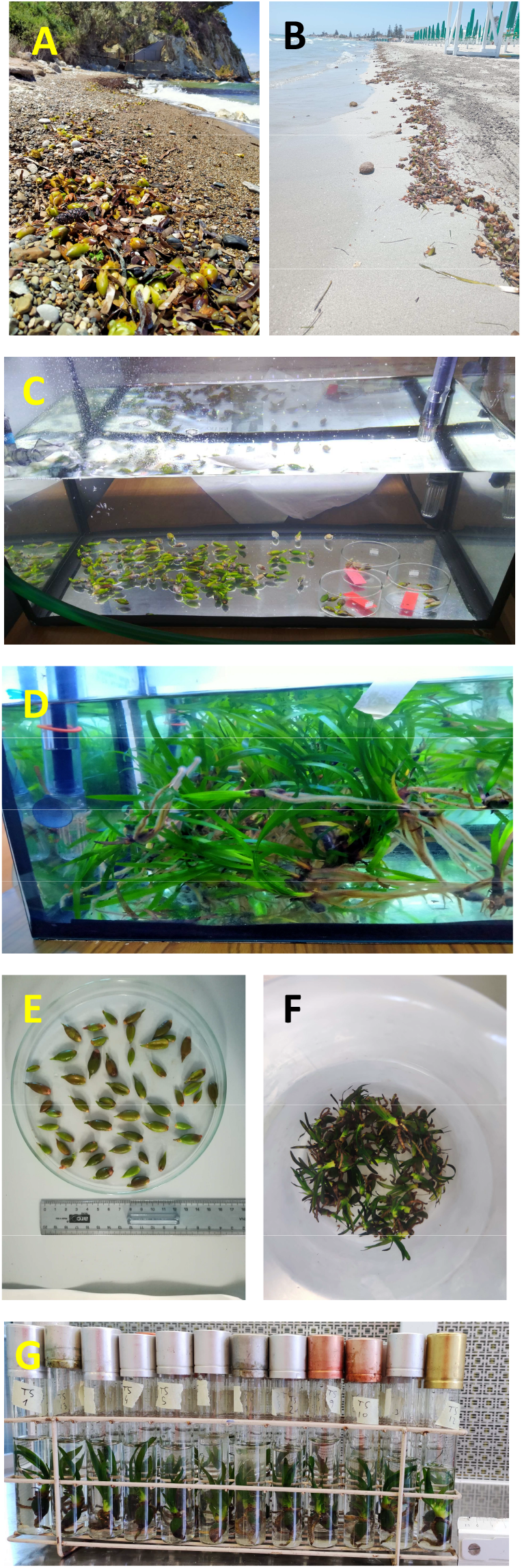
Stranded seeds and fruits at the collection events on the beaches of Trabia (A); and Mazara del Vallo (B). Fresh seeds in aquarium (C); and after nine months (D). Fresh T0 seeds ready to be stored (E). T2 seedlings stored in a beaker (F); or in individual tubes (G).

## Acknowledgements

This work was supported by the project Marine Hazard, PON03PE_00203_1 (Ministry of Education, University and Research, MUR, Italy).

## Author contributions

D.P, F.C. and R.D.M. conceived the ideas; F.C. and R.D.M. provided the funding; R.D.M designed the methodology; R.D.M. collected the seeds; A.S. and P.S., performed the experiments; R.D.M. analyzed the data; R.D.M. wrote the manuscript. All authors contributed critically to the drafts and gave final approval for publication.

## References

Alagna, A., D’Anna, G., Musco, L., Vega Fernández, T., Gresta, M., Pierozzi, N., & Badalamenti, F. (2019). Taking advantage of seagrass recovery potential to develop novel and effective meadow rehabilitation methods. Marine Pollution Bulletin, 149, 110578. 10.1016/J.MARPOLBUL.2019.110578

Alagna, A., Fernández, T. V, Anna, G. D., Magliola, C., Mazzola, S., & Badalamenti, F. (2015). Assessing Posidonia oceanica Seedling Substrate Preference: An Experimental Determination of Seedling Anchorage Success in Rocky vs. Sandy Substrates. Sandy Substrates. PLoS ONE, 10(4), 125321. 10.1371/journal.pone.0125321

Alagna, A., Giacalone, V. M., Zenone, A., Martinez, M., D’Anna, G., Buffa, G., Cavalca, C. J., Poli, A., Varese, G. C., Prigione, V. P., & Badalamenti, F. (2024). Tannins and copper sulphate as antimicrobial agents to prevent contamination of Posidonia oceanica seedling culture for restoration purposes. Frontiers in Plant Science, 15, 1433358. 10.3389/FPLS.2024.1433358/BIBTEX

Alagna, A., Zenone, A., & Badalamenti, F. (2020). The perfect microsite: How to maximize Posidonia oceanica seedling settlement success for restoration purposes using ecological knowledge. Marine Environmental Research, 161, 104846. 10.1016/J.MARENVRES.2019.104846

Balestri, E., Piazzi, L., & Cinelli, F. (1998). In vitro germination and seedling development of Posidonia oceanica. Aquatic Botany, 60(1), 83–93. 10.1016/S0304-3770(97)00017-X

Balestri, E., Vallerini, F., & Lardicci, C. (2017). Recruitment and patch establishment by seed in the seagrass Posidonia oceanica: Importance and conservation implications. Frontiers in Plant Science, 8, 1067. 10.3389/FPLS.2017.01067/BIBTEX

Belzunce, M., Navarro, R. M., & Rapoport, H. F. (2008). Posidonia oceanica seeds from drift origin: Viability, germination and early plantlet development. Botanica Marina, 51(1), 1–9. 10.1515/BOT.2008.005/MACHINEREADABLECITATION/RIS

Bradamante, G., Scheid, O. M., & Incarbone, M. (2021). Under siege: virus control in plant meristems and progeny. 10.1093/plcell/koab140

Buia, M. C., & Mazzella, L. (1991). Reproductive phenology of the Mediterranean seagrasses Posidonia oceanica (L.) Delile, Cymodocea nodosa (Ucria) Aschers., and Zostera noltii Hornem. Aquatic Botany, 40(4), 343–362. 10.1016/0304-3770(91)90080-O

Calvo, S., Tomasello, A., Di Maida, G., Pirrotta, M., Buia, M. C., Cinelli, F., Cormaci, M., Furnari, G., Giaccone, G., Luzzu, F., Mazzola, A., Orestano, C., Procaccini, G., Sarà, G., Scannavino, A., & Vizzini, S. (2010). Seagrasses along the Sicilian coasts. Https://Doi.Org/10.1080/02757541003636374, 26(SUPPL. 1), 249–266. 10.1080/02757541003636374

Celdran, D. (2013). Seed photosynthesis enhances Posidonia oceanica seedling growth Seed collection. Ecosphere, 4(December), 1–11.

Diaz-Almela, E., Marbà, N., & Duarte, C. M. (2007). Consequences of Mediterranean warming events in seagrass (Posidonia oceanica) flowering records. Global Change Biology, 13(1), 224–235. 10.1111/J.1365-2486.2006.01260.X

Escandell-Westcott, A., Riera, R., & Hernández-Muñoz, N. (2023). Posidonia oceanica restoration review: Factors affecting seedlings. Journal of Sea Research, 191, 102337. 10.1016/j.seares.2023.102337

Goggin, D. E., Steadman, K. J., Emery, R. J. N., Farrow, S. C., Benech-Arnold, R. L., & Powles, S. B. (2009). ABA inhibits germination but not dormancy release in mature imbibed seeds of Lolium rigidum Gaud. Journal of Experimental Botany, 60(12), 3387–3396. 10.1093/JXB/ERP175

Infantes, E., Orfila, A., Bouma, T. J., Simarro, G., & Terrados, J. (n.d.). Posidonia oceanica and Cymodocea nodosa seedling tolerance to wave exposure. 10.4319/lo.2011.56.6.2223

Khan, A. A. (1994). Induction of Dormancy in Nondormant Seeds. Journal of the American Society for Horticultural Science, 119(3), 408–413.

Marbà, N., & Duarte, C. M. (1998). Rhizome elongation and seagrass clonal growth. Marine Ecology Progress Series, 174, 269–280. 10.3354/MEPS174269

Nakabayashi, K., Bartsch, M., Xiang, Y., Miatton, E., Pellengahr, S., Yano, R., Seo, M., & Soppe, W. J. J. (2012). The Time Required for Dormancy Release in Arabidopsis Is Determined by DELAY OF GERMINATION1 Protein Levels in Freshly Harvested Seeds. The Plant Cell, 24(7), 2826–2838. 10.1105/TPC.112.100214

Orth, R. J., Harwell, M. C., Bailey, E. M., Orth, R. J.;, Harwell, M.;, Bailey, E.;, Bartholomew, A.;, Jawad, J.;, Lombana, A.;, Moore, K.;, & Rhode, J.; (2000). A review of issues in seagrass seed dormancy and germination: implications for conservation and restoration. MARINE ECOLOGY PROGRESS SERIES, 200, 277. 10.3354/meps200277

Scanu, S., Piazzolla, D., Bonamano, S., Penna, M., Piermattei, V., Madonia, A., Frattarelli, F. M., Mellini, S., Dolce, T., Valentini, R., Coppini, G., Fersini, G., & Marcelli, M. (2022). Economic Evaluation of Posidonia oceanica Ecosystem Services along the Italian Coast. Sustainability 2022, Vol. 14, Page 489, 14(1), 489. 10.3390/SU14010489

Steinbach, H. S., Benech-Arnold, R. L., & Sánchez, R. A. (1997). Hormonal Regulation of Dormancy in Developing Sorghum Seeds. Plant Physiology, 113(1), 149–154. 10.1104/PP.113.1.149

Sullivan, B. K., Trevathan-Tackett, S. M., Neuhauser, S., & Govers, L. L. (2018). Review: Host-pathogen dynamics of seagrass diseases under future global change. Marine Pollution Bulletin, 134, 75–88. 10.1016/J.MARPOLBUL.2017.09.030

Sutera, A., Bonaviri, C., Spinelli, P., Carimi, F., & De Michele, R. (2024). Fruit encasing preserves the dispersal potential and viability of stranded Posidonia oceanica seeds. Scientific Reports 2024 14:1, 14(1), 1–8. 10.1038/s41598-024-56536-x

Telesca, L., Belluscio, A., Criscoli, A., Ardizzone, G., Apostolaki, E. T., Fraschetti, S., Gristina, M., Knittweis, L., Martin, C. S., Pergent, G., Alagna, A., Badalamenti, F., Garofalo, G., Gerakaris, V., Louise Pace, M., Pergent-Martini, C., & Salomidi, M. (2015). Seagrass meadows (Posidonia oceanica) distribution and trajectories of change. Scientific Reports 2015 5:1, 5(1), 1–14. 10.1038/srep12505

Zenone, A., Badalamenti, F., Alagna, A., Gorb, S. N., & Infantes, E. (2022). Assessing Tolerance to the Hydrodynamic Exposure of Posidonia oceanica Seedlings Anchored to Rocky Substrates. Frontiers in Marine Science, 8, 788448. 10.3389/FMARS.2021.788448/BIBTEX

Zenone, A., Giacalone, V. M., Martinez, M., Pipitone, C., Alagna, A., Infantes, E., D’Anna, G., & Badalamenti, F. (2025). Stitching up Posidonia oceanica (L.) Delile anchorage scars using beach-cast seeds: Results of a six-year study. Biological Conservation, 303, 111032. 10.1016/J.BIOCON.2025.111032

